# Network of small no-take marine reserves reveal greater abundance and body size of fisheries target species

**DOI:** 10.1101/422147

**Authors:** Fernanda A. Rolim, Tim Langlois, Pedro F. C. Rodrigues, Todd Bond, Fábio S. Motta, Leonardo M. Neves, Otto B. F. Gadig

## Abstract

No-take marine reserves (NTRs), i.e. areas with total fishing restrictions, have been established worldwide aiming to provide biodiversity and ecosystem conservation. Brazil has established NTRs, but currently lacks scientific knowledge and understanding of their ecological role, particularly regarding rocky reefs in subtropical regions. Consequently, this study aimed to contrast NTR, from the 30 year old Tupinambás Ecological Station, and comparable fished sites across a coastal biogeographic gradient to investigate the effect of fishing and habitat variability on the abundance and body-size of rocky reef fish. We used Baited Remote Underwater stereo-Video (stereo-BRUVs) and Diver Operated stereo-Video (stereo-DOVs) systems to simultaneously sample reef fish and habitat. Model selection and results identified habitat and biogeographic variables, such as distance from shore, as important predictor variables, explaining several aspects of the fish assemblage. The effect of protection was important in determining the abundance and body-size of targeted species, in particular for epinephelids and carangids. Conversely, species richness was correlated with habitat complexity and not with protection status. This is the first study to publish data using these novel survey methods in the Southeastern Atlantic and demonstrate their utility, in combination with an established network of NTR’s, to provide benchmarks for conservation and fisheries management.

## Introduction

No-take marine reserves (NTRs) have been established worldwide as a management strategy mostly aiming to recover marine biodiversity from effects of fisheries. It is well documented that these areas can provide refuge to marine life, increasing local abundance, species richness, body size and the reproductive capacity of fish [1–4].

Networks of NTRs can be used to investigate effects of fishing across biogeographic gradients, with the aim of estimating benchmarks for conservation and fisheries management. Besides facing direct fisheries effects from biomass removal, NTRs have also highlighted indirect effects, where the removal of large carnivores in areas without protection can result in a higher abundance of prey species, leading to a trophic reorganization [5–7].

It is also well documented that fish assemblage structure will vary with physical, chemical and biological factors across biogeographic and habitat gradients through space [8– 10]. In particular, distance from the coast and topographic complexity have shown to increase species richness, abundance and biomass of reef fish [11–16]. It is therefore important for any investigation of the effects of fishing to be able to control for these covariates across any inside versus outside NTR comparison.

Brazil has 8500 km of coastline and a territorial sea that, together with the Exclusive Economic Zone, encompasses 4 million km^2^. Of this area, 26.4% is protected by 177 marine protected areas (MPAs), of which 73 are NTR representing 3.3% of the country’s marine waters [17]. However, the few studies available about the effects of Brazilian NTRs on fish assemblage are concentrated in the northern part [18], southern part [19, 20] or offshore islands [21, 22], with a lack a studies in the transition zones and small coastal NTR networks.

Coastal habitat on the northern coast of Brazil (north of 19°S) are dominated by coral reefs, whereas in southern regions (between 19-28°S) they are typified by rocky reefs. In general, the Brazilian province concentrates high number of endemic species and biomass of organisms [23–26]. In the transition zone between tropical and subtropical-temperate environments (20°S to 23°S), the mixture of habitat types results in some of the highest species diversity of benthic [27] and reef fish species recorded in Brazil [23]. These transitional reefs are biologically rich and complex environments where it is vitally important to use the established network of NTR’s, in the region, to increase our ecological understanding and provide benchmarks for conservation and fisheries management.

Historically, NTRs in Southwestern Atlantic have been assessed using underwater visual census (UVC) (e.g. [18, 19, 21, 22]). However, biases involving interobserver variability, underrepresentation of species targeted by fisheries, as well as inaccuracy of species identification and size estimates are likely to occur [28, 29]. In order to overcome this issue, the use of videos to collect data has been increasingly adopted [30, 31], especially since rapid advancements in video technology and accessibility to cheaper and higher quality equipment. Importantly, methods using such technologies create a permanent record allowing fish identification to be confirmed by experts and revisited when necessary.

Baited Remote Underwater stereo-Video (stereo-BRUV) and Diver Operated stereo-Video (stereo-DOV) are being widely employed to assess diverse aspects of fish assemblages [32–35]. Stereo-video techniques provide accurate body-size and range measurements of individuals from the three-dimensional calibration of imagery (Harvey and Shortis, 1996). Stereo-BRUV have been found to sample a wide range of species without precluding estimates of herbivorous species [36] and can be applied across a wide variety of habitats and depths [34]. Also, as a remote sensing technique, it detects large and mobile animals which usually avoid divers and active fishing gears [32, 33], but has a range of acknowledge biases and limitations (see Langlois et al. [37]). Conversely, the presence of a diver may impact the abundance of fish recorded using stereo-DOVs [33, 38], suggesting that a combination of methods more effectively samples the fish assemblage [33, 39].

In expanding our knowledge of ichthyofauna in the Southeastern Atlantic, it is important that non-destructive sampling methods are adopted that enable robust estimates of fisheries species inside and outside NTRs. This study is the first assessing fish assemblages using stereo-BRUVs and stereo-DOVs in Southwestern Atlantic and Brazil, and aims to to generate robust data for conservation and fisheries management in the region. We aim to investigate the response of the fish assemblage to environmental and habitat variables, as well as the effect of fishing using comparison with NTRs. We hypothesize that: (1) abundance and body-size of targeted fish groups will be greater inside NTRs; whereas (3) non-target fish abundance and species richness will be explained better by habitat and biogeographic variables.

## Material and methods

### Study site

The Ecological Station (ESEC) of Tupinambás is a no-take reserve located on the northern coast of São Paulo State, Brazil, Southeastern Atlantic. The ESEC was established in 1987 [40] and is divided into two sectors; sector I is in the archipelago of Alcatrazes located approximately 43 km from of São Sebastião, São Paulo. Sector I has six key management sites, each with a 1 km buffer of protected area. Two sets of two sites are close enough to overlap, creating four primary areas of protection (Fig 1). Sector II protects Palmas Island including two nearby slab reefs (Palmas Slab and Forno Slab) and Cabras Island, located 5.7 km and 3.6 km respectively from the coast of Ubatuba, São Paulo. An ESEC is a fully protected area (corresponding to IUCN Category Ia), allowing only scientific research.

**Fig 1.**
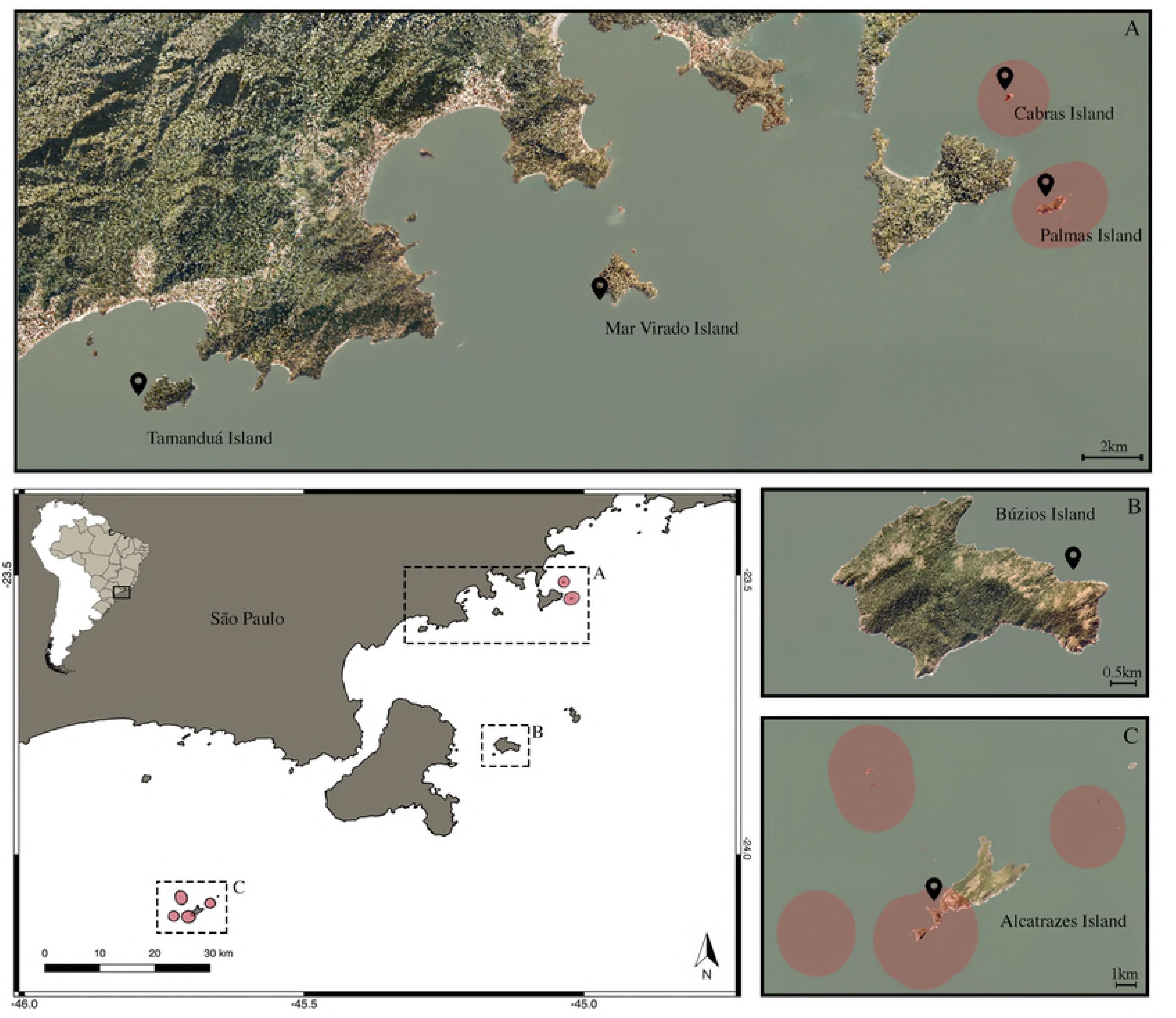
Map of the study area on the north coast of São Paulo state with the no-take marine reserve Tupinambás Ecological Station in red. The control islands, where fishing activity is permitted (Tamanduá, Mar Virado and Búzios) is also displayed (A, B). ESEC areas in detail in the Islands of Cabras and Palmas (A) and Alcatrazes Archipelago (C) with the sample sites represented by the black spots. Source of pictures: Emplasa [41] and CFA/DPM [42].

The open-fishing areas used to test the effects of protection on fish assemblage included Búzios, Mar Virado and Tamanduá islands. These islands are part of a multiple use MPA established in 2008 (Environmental Protection Area: EPA - corresponding to IUCN category V). The islands are located 34 km, 2 km, and 0.5 km respectively from the mainland. Small scale fishing is permitted around Mar Virado and Tamanduá island, but no industrial fishing that uses pair trawling, driftnet vessels above 20 gross tonnage (GT) or trawling vessels up to 10 GT is permitted. However, only pair trawlers are excluded from fishing in Búzios island.

### Sampling

Samples were collected using stereo-DOVs and stereo-BRUVs. Both equipment types are comprised of a metal base bar with two underwater SeaGIS housing (www.seagis.com.au), each with a digital video cameras inside. Housings are positioned approximately 700 mm apart, each inwardly converged at 8 degrees towards the center. Stereo-BRUVs were deployed from a boat with a connected roped and surface float and left on the seafloor to record. The camera base bar was enclosed within a stainless steel frame and a bait cage with 800 g of mashed sardine (*Sardinella brasiliensis*) was positioned at the end of a bait arm approximately 1.5 m away from the cameras. Oily bait provide greater sampling efficiency [43, 44] odor plume dissipates more efficiently. Stereo-DOVs used the same camera base bar setup with the additional of handle allowing divers to swim a transect. These standard survey methods have been developed and used by many authors worldwide [32, 34, 45].

Stereo-BRUV and stereo-DOV sampling was undertaken bimonthly at each island for a year (2016 - March, May, July, October, November; and 2017 - January). Each expedition was approximately 4-6 days long, covering all six islands. Due to the small size of islands and to maintain independence among samples, (suggested minimum distance between replicates is at least 250 m [36]), only two stereo-BRUV samples were collected on the leeward side of the islands, totaling 12 stereo-BRUVs at each island after six expeditions. Each stereo-BRUV was deployed at the interface of the rocky reef with the sandy bottom and left to soak for 90 minutes. Water depth ranged from 2-17 m depending on the location of the rock-sand interface at each island and the average water depth sampled was 8.3±3.6 m.

Stereo-DOV transects were 25 m long and 5 m wide, and swum at the interface of the rocky shore with the sandy bottom, as well as at the shallow zone above the reef. Due to the small size of the islands, sampling was restricted to three transects at the rock-sand interface and three in the shallow zone on each island ateach expedition, totaling 36 transects per island at the end of six expeditions. Transects were swum in two passes, the first pass filmed fish in the water column, the second returning pass filmed benthic and more cryptic fish species. The interface zone presented an average depth of 8.9±3.8 m and the shallow zone 4.2±1.9 m.

### Video analysis

#### Fish assemblage

Stereo-video systems were calibrated using the CAL software and video analysis was carried out in the EventMeasure software (www.seagis.com.au). Fish were identified to the finest taxonomic level possible, counted and measured if they were within 7m of the stereo-BRUVs and 5m for stereo-DOVs.

The relative abundance of each species filmed on stereo-BRUVs was recorded as MaxN, defined as the maximum number of individuals of the same species recorded in a single frame from the left camera. This is a conservative approach in order to avoid counting and measuring the same individual more than once. The fork length of individual fish contributing to a species’ MaxN was measured when the fish was straight and no more than 45 degrees perpendicular to the cameras. In the stereo-DOV, all fish filmed on the left camera were counted and measured using the same rules. These data are stored on GlobalArchive [46].

Biomass was calculated for all species using measured fish lengths and length-weight relationship referenced in the FishBase database [47]. If equations for fork length of a species were not available, length-length conversions were used if available. Biomass of species without length-weight information was calculated using equations from a similar species from the same family.

Fish species were classified by broad functional groups based on diet, using information available in the literature [48, 49] and FishBase [47]. Groups included: carnivores, piscivores, planktivores, roving herbivores, territorial herbivores, omnivores, sessile invertivores, mobile invertivores. Piscivores were pooled with carnivores because there was not enough individuals for statistical analysis.

Species were categorized in target and non-target for fisheries in the region according to the literature [16, 50–52]. Four families (Epinephelidae, Kyphosidae, Scaridae and Carangidae) identified as abundant or frequent and also targeted by fisheries were selected for analysis.

#### Habitat characteristics

Habitat classification and complexity (mean relief) was analysed using a single high definition image each stereo-BRUV deployment and three single frames of each stereo-DOV transect separated by approximately 8 m. This method is shown to be effective to determine reefs structural complexity [53–55]. Images were analyzed in TransectMeasure software (www.seagis.com.au) using a standardised broad habitat classification scheme based on CATAMI [56] to classify benthic composition and based on [57] to classify relief characteristics (Table 2). Each image was divided into a 5 × 4 grid and the dominant habitat type of each square was recorded. An additional category, ‘reef’, was formed by pooling macroalgae, stony coral, rock and zoanthids, and is based on the similar broad structure these environments present.

**Table 1.**
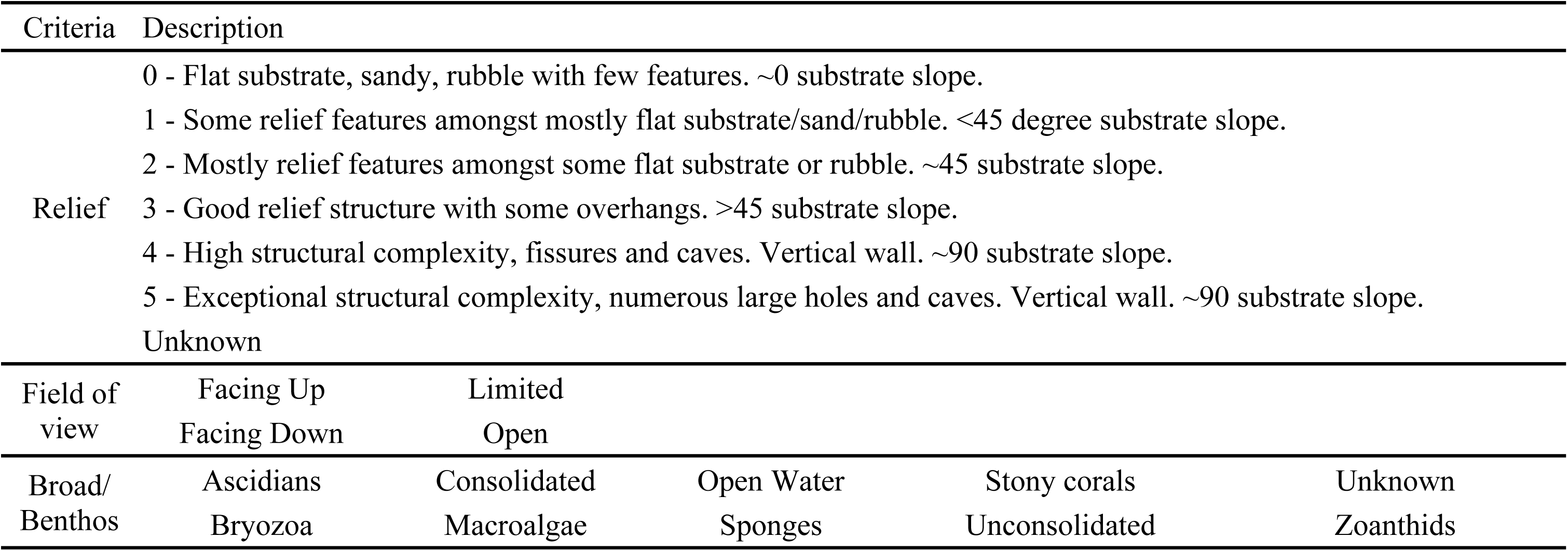
Habitat classification based on broad CATAMI Classification scheme [56] and on [57], used in Baited Remote Underwater stereo-Videos and Diver Operated stereo-Videos images.

**Table 2.**
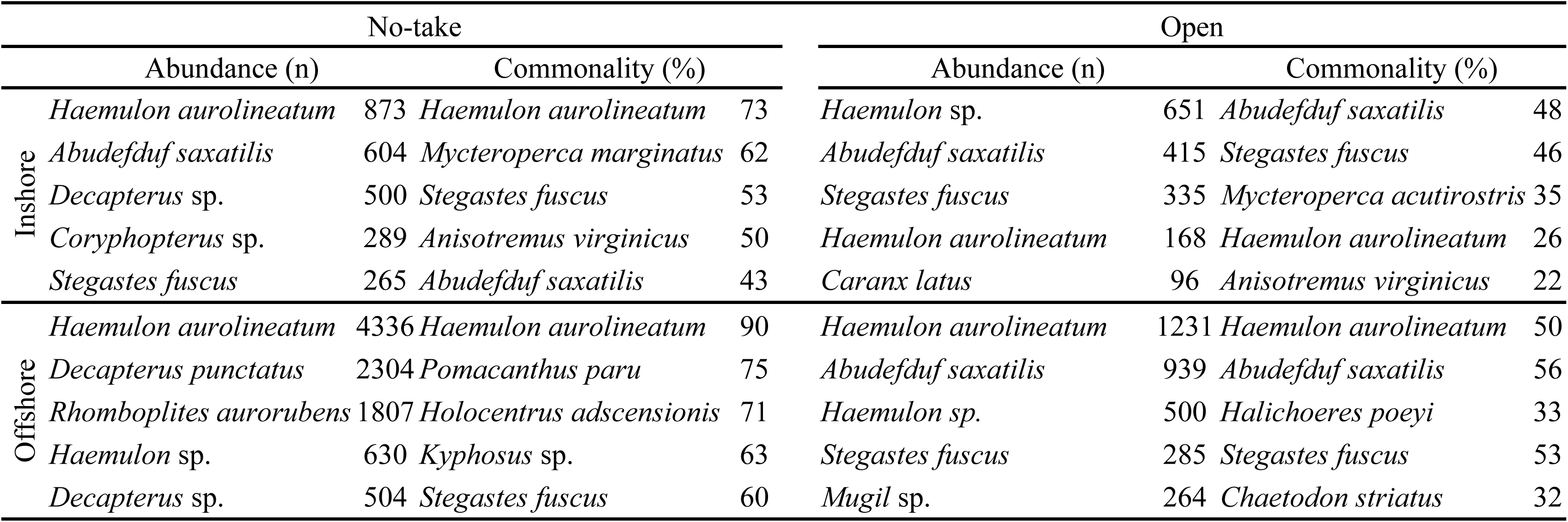
Top five most abundant and common species (% of samples a species was observed) in no-take and open to fisheries areas in inshore and offshore regions.

#### Environmental variables

Environmental variables were recorded at each sampling event. Temperature and salinity were measured using a Castaway® CTD (Conductivity, Temperature and Depth) and an average temperature and salinity value was calculated from values recorded at the BRUV or dive depth, and 1 m above and below this. Visibility was estimated using a Secchi disk.

### Data analysis

The influence of habitat characteristics and environmental variables on fish assemblage richness, abundance and biomass was investigated using Generalized Additive Mixed Models (GAMM) [58, 59] and a full-subsets multiple regression approach based on the function described by [60]. GAMMs use smoothing splines to estimate non-parametric additive functions, allowing for overdispersion and correlation in the data [58], which may arise in studies like this.

Models were fitted to untransformed overall abundance, richness and biomass data, as well as to abundance by functional group and by families. Models for biomass by functional group and by family were also determined, however, as the same trends were found, we decided to report results on abundance only. A prior selection of the predictor variables was made based on their coverage and on the high collinearity between them (Pearson correlation coefficient r > 0.8). As a result, Reef, Rock and Mean relief remained as continuous variables for the analysis. Null variables of the random model included Month, Method, Depth and Visibility, and fixed factors included Distance to shore (two levels: inshore and offshore) and Protection (two levels: no-take and open). Continuous predictor variables were square root transformed to reduce dispersion of data.

Model selection for each response variable was based on the second-order variant of Akaike’s Information Criterion suited for small samples (AIC_*C*_) [61] and on AIC_*c*_ weights (ωAIC_*c*_). The best model was the most parsimonious one (with the fewest variables) within two AIC_*c*_ units of the lowest AIC_*c*_ value (ΔAIC_*c*_<2) [62]. Because the effect of protection status, and any interactions, were relevant to the primary hypothesis of this study, models that were within two AICc units of the model with the lowest AIC_*c*_ and included protection status, were therefore preferentially investigated (‘hypothesis model’). Selected models had their shape and effective degrees of freedom (EDF) examined to ensure they did not overfit the data.

The distribution of fish lengths for key families were compared inside and outside NTRs using Mann-Whitney *U* test, considering a significant difference as p-values below 0.05. All analyses were performed using R Language for Statistical Computing [63], with the packages gamm4 [64], mgcv [65], MuMIn [66], doParallel [67] and dplyr [68].

## Results

A total of 23,505 individuals were observed belonging to 126 species of 44 families (list in S1 Table). Large schools (>100) of sardines (Clupeidae), mullets (*Mugil* spp.), young mackerels (*Decapterus* spp.), young vermilion snapper (*Rhomboplites aurorubens*) and young grunts (Haemulidae) were excluded from statistical analysis in order to reduce dispersion of data and highlight effects. Not considering these schools, the most abundant and frequent families were grunts (Haemulidae), damselfishes (Pomacentridae), jacks (Carangidae) and snappers (Lutjanidae). The most abundant species were, in this order, *Haemulon aurolineatum* (28.1%), *Abudefduf saxatilis* (10.1%), *Stegastes fuscus* (4.7%), *Coryphopterus sp.* (2.3%) and *Holocentrus adscensionis* (1.3%). And the most frequent species were *Haemulon aurolineatum* (64.2%), *Stegastes fuscus* (54.2%), *Abudefduf saxatilis* (49.6%), *Anisotremus virginicus* (38.9%) and *Mycteroperca marginatus* (35.1%).

*Haemulon aurolineatum* was listed as a highly common and abundant species in inshore and offshore in both no-take and fished areas (Table 2). *Stegastes fuscus* and *Abudefduf saxatilis* was also highly recorded as abundant and frequent, except for the abundance in offshore no-take areas, which was mainly represented by schools of grunts, mackerels and vermilion snappers (Table 2).

Nine species are endemic to the Brazilian Province [69–71] and fourteen species are considered threatened (vulnerable/endangered) or near threatened, by the International Union for Conservation of Nature (IUCN) Red List and the Brazilian legislation [72] (detailed list in S1 Table).

The most parsimonious model for total richness included distance to shore and mean relief, whereas for both total abundance and biomass the selected models included protection status and distance to shore (Table 3, Figs 2 and 3). The model for overall abundance was selected based on the primary hypothesis of interest, and was within 2AIC of the top model, but it is interesting to note that mean relief was highly important (Fig 2) and present in the most parsimonious model.

**Table 3.** Top generalised additive mixed models (GAMMs) to predict different aspects of fish assemblage. ΔAIC_*c*_= Difference between lowest reported corrected Akaike Information Criterion; ΔBIC= Bayesian Information Criterion; ωAIC_*c*_= AIC_*c*_ weights; ωBIC= BIC weights; R^2^= variance explained; EDF= effective degrees of freedom. Model selection was based on the most parsimonious model within two units of the lowest AIC_*c*_ which has the fewest variables.

| Dependent variables | Best models | $\Delta AIC_c$ | $\Delta BIC$ | $\omega AIC_c$ | $\omega BIC$ | $R^2$ | EDF |
| --- | --- | --- | --- | --- | --- | --- | --- |
| **Overall** |  |  |  |  |  |  |  |
| Richness | Distance to shore + Mean relief.by.Distance to shore | 0.00 | 0.00 | 1.00 | 0.89 | 0.57 | 18.75 |
| Abundance | Mean relief + Distance to shore | 0.49 | 0.00 | 0.32 | 0.68 | 0.30 | 8.53 |
|  | Protection + Distance to shore | 0.00 | 1.57 | 0.41 | 0.31 | 0.34 | 8.76 |
|  | Distance to shore + Mean relief.by.Distance to shore | 0.89 | 7.52 | 0.26 | 0.02 | 0.30 | 10.33 |
| Biomass | Protection + Distance to shore | 0.00 | 0.00 | 1.00 | 1.00 | 0.27 | 16.50 |
| **Importance to fisheries** |  |  |  |  |  |  |  |
| Non-target species richness | Distance to shore + Mean relief.by.Distance to shore | 0.00 | 0.00 | 1.00 | 1.00 | 0.47 | 14.53 |
| Non-target species abundance | Distance to shore + Mean relief.by.Distance to shore | 0.00 | 4.83 | 0.90 | 0.08 | 0.39 | 8.54 |
| Target species richness | Mean relief + Distance to shore | 0.00 | 0.00 | 0.61 | 0.89 | 0.56 | 18.42 |
| Target species abundance | Protection + Distance to shore | 0.00 | 0.00 | 1.00 | 1.00 | 0.25 | 13.55 |
| **Abundance by functional group** |  |  |  |  |  |  |  |
| Carnivores/Piscivores | Protection + Mean relief.by.Protection | 0.00 | 6.50 | 0.86 | 0.02 | 0.30 | 17.28 |
| Mobile invertivores | Protection + Distance to shore | 0.00 | 0.00 | 1.00 | 1.00 | 0.23 | 10.21 |
| Sessile invertivores | Protection | 1.43 | 0.00 | 0.15 | 0.40 | 0.13 | 7.31 |
|  | Protection + Reef.by.Protection | 0.17 | 4.73 | 0.28 | 0.04 | 0.16 | 9.16 |
|  | Protection + Mean relief.by.Protection | 0.00 | 12.54 | 0.31 | 0.00 | 0.11 | 10.92 |
| Omnivores | Mean relief + Distance to shore | 0.00 | 0.00 | 0.90 | 0.96 | 0.31 | 13.38 |
| Planktivores | Protection + Mean relief.by.Protection | 0.00 | 3.54 | 0.92 | 0.14 | 0.03 | 8.81 |
| Roving herbivores | Mean relief + Distance to shore | 0.00 | 0.00 | 0.44 | 0.48 | 0.17 | 6.95 |
|  | Distance to shore + Mean relief.by.Distance to shore | 0.47 | 17.10 | 0.35 | 0.00 | 0.18 | 9.22 |
| Territorial herbivores | Protection + Mean relief.by.Protection | 0.00 | 0.00 | 0.89 | 0.93 | 0.40 | 14.66 |
| **Abundance by family** |  |  |  |  |  |  |  |
| Epinephelidae | Protection + Reef.by.Protection | 0.00 | 0.00 | 1.00 | 1.00 | 0.27 | 19.49 |
| Kyphosidae | Distance to shore + Reef.by.Distance to shore | 0.00 | 0.00 | 0.53 | 0.42 | 0.15 | 17.92 |
|  | Distance to shore + Rock.by.Distance to shore | 0.30 | 0.58 | 0.46 | 0.31 | 0.16 | 17.94 |
| Scaridae | Protection + Distance to shore | 0.00 | 0.00 | 1.00 | 1.00 | 0.10 | 15.14 |
| Carangidae | Protection + Reef.by.Protection | 0.00 | 0.00 | 1.00 | 1.00 | 0.22 | 20.22 |

**Fig 2.**
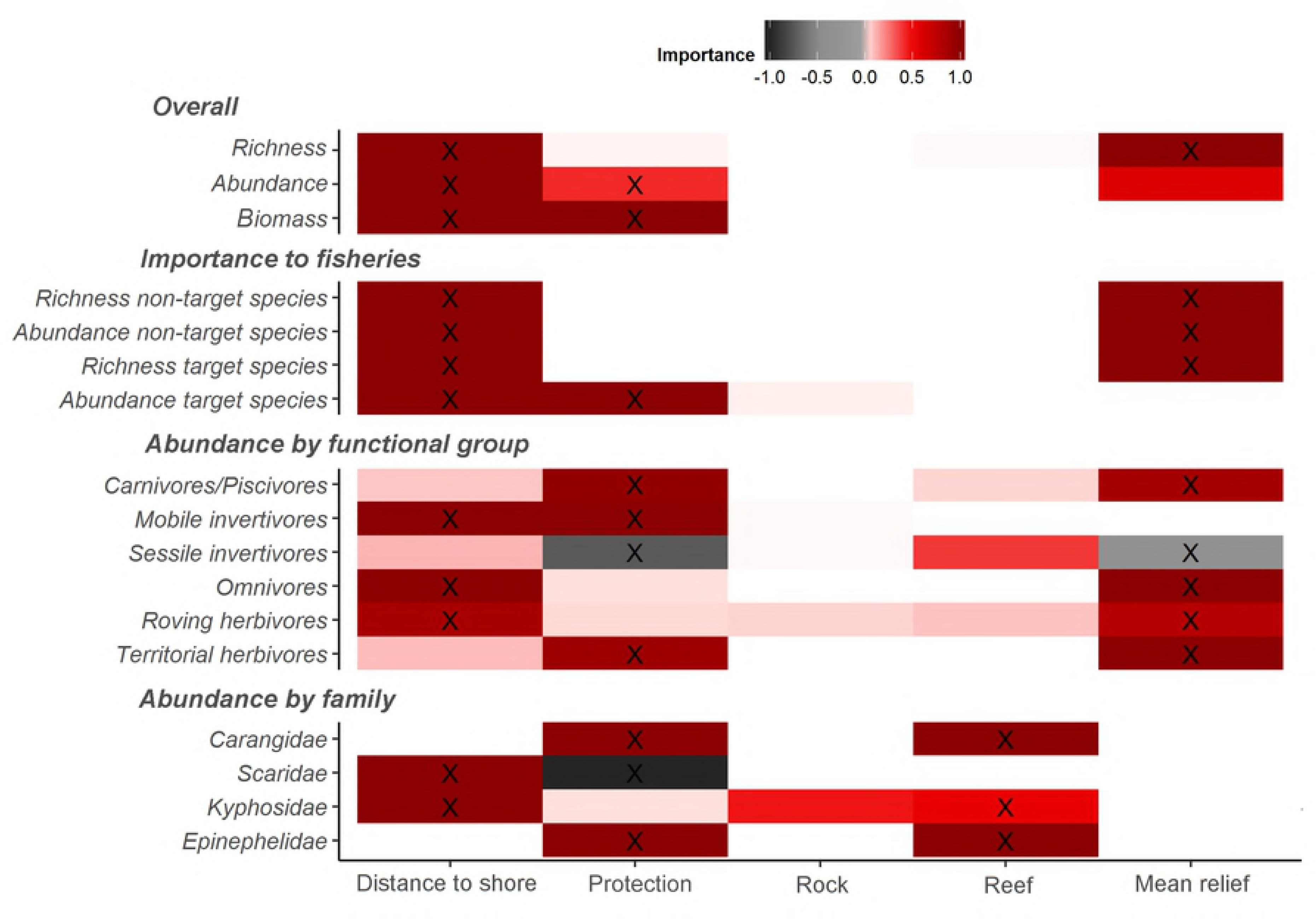
Variable importance scores from full-subset generalised additive mixed models analysis, with >10% variance explained shown. X= Predictor variables within the most parsimonious model for each response variable (see Table 1).

**Fig 3.**
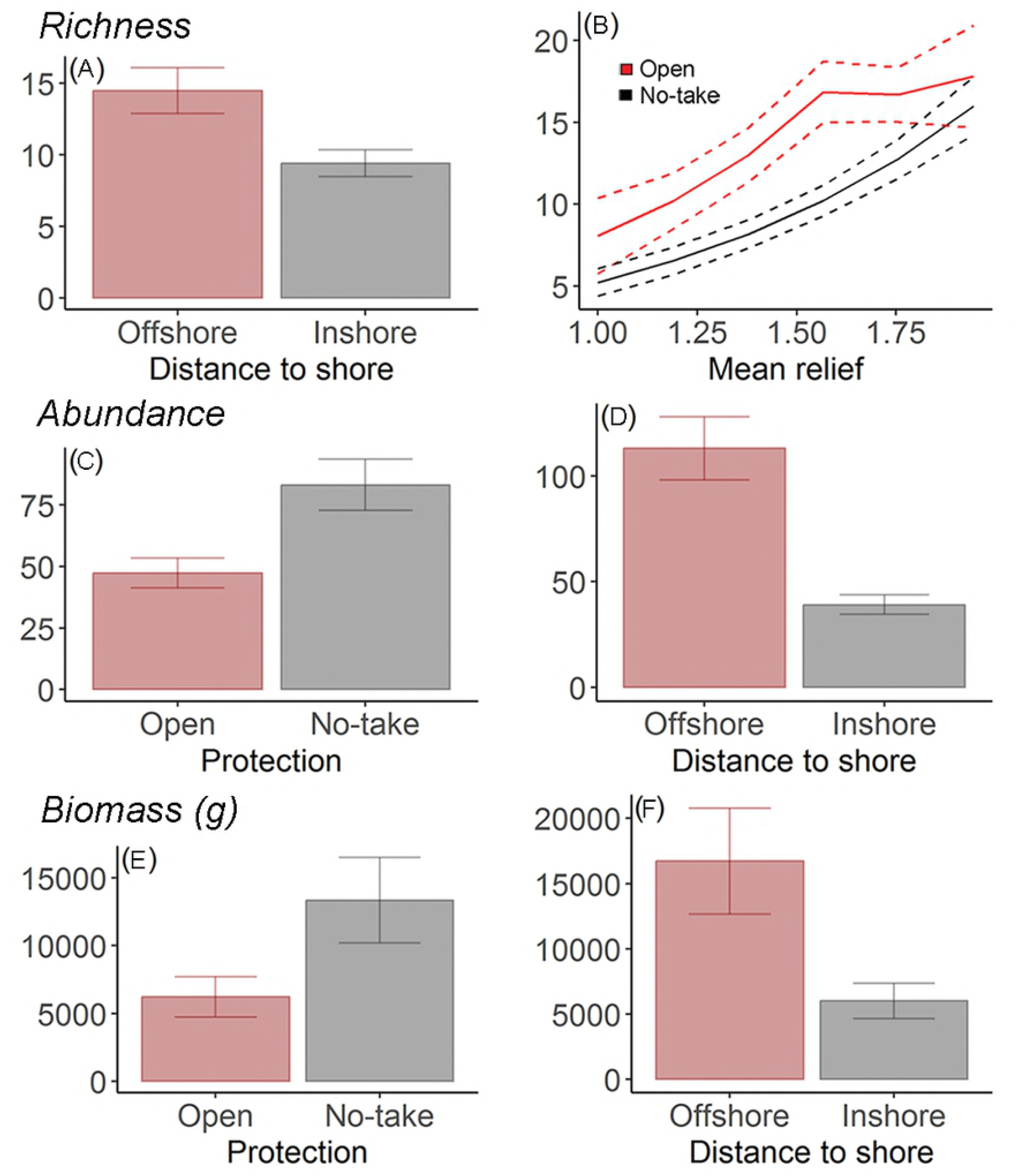
Plots of the most parsimonious model for (A,B) species richness, (C,D) total abundance and (E,F) total biomass. The dotted line represents 95% confidence interval.

Higher richness of target and non-target species and greater abundance of non-target species were best predicted by increased distance to shore and mean relief, whilst the most parsimonious models for the abundance of target species indicated they were likely to increase with protection and distance to shore (Fig 4). Concerning abundance by functional groups, the most parsimonious models for carnivores/piscivores, planktivores and territorial herbivores all included a positive relationship with protection and mean relief. However, abundance of sessile invertivores was found to be negatively correlated with protection status and mean relief. For mobile invertivores, the abundance is likely to increase with protection and distance to shore. The number of herbivores and omnivores was higher in areas further from shore and also on structurally complex reefs (Fig 5).

**Fig 4.**
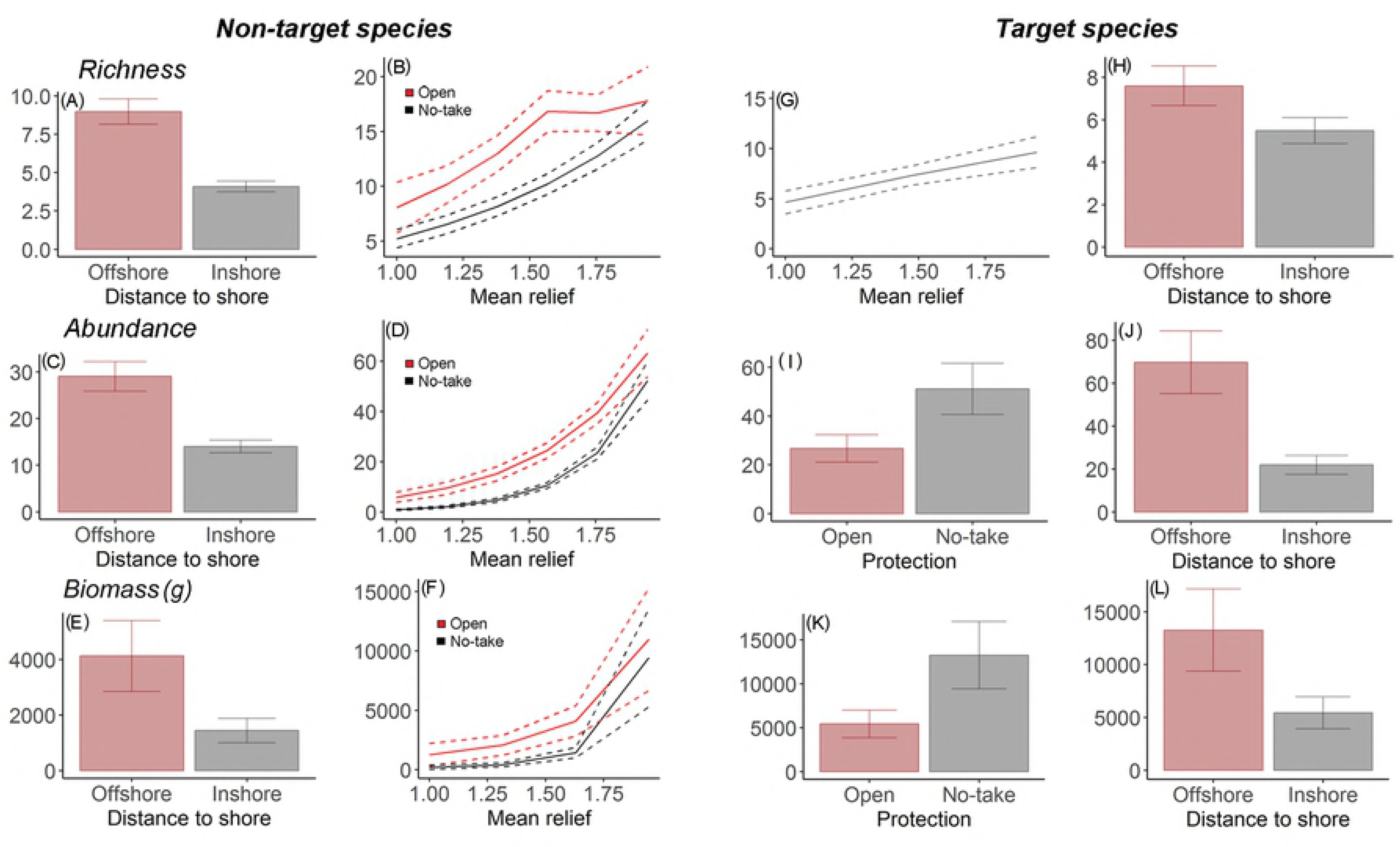
Plots of the most parsimonious model for non-target species (A,B) richness, (C,D) abundance, (E,F) biomass. And for target species (G,H) richness, (I,J) abundance, (K,L) biomass. The dotted line represents 95% confidence interval.

**Fig 5.**
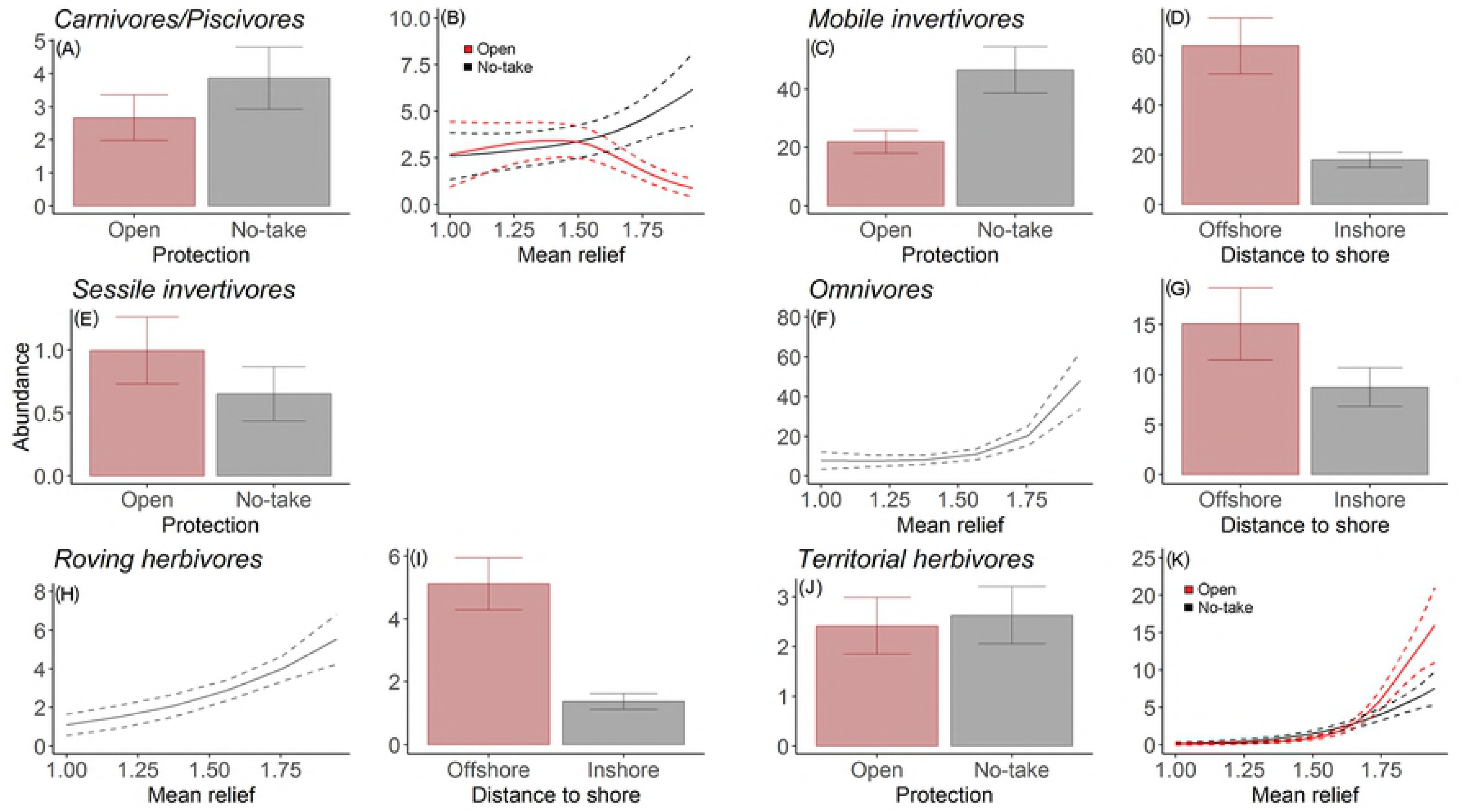
Plots of the most parsimonious model for (A,B) Carnivores/piscivores, (C,D) Mobile invertivores, (E) Sessile invertivores, (F,G) Omnivores,(H,I) Roving herbivores, (J,K) Territorial herbivores. The dotted line represents 95% confidence interval.

Targeted families Carangidae and Epinephelidae increased with protection and presence of reef, whilst kyphosids were found in greater abundance in areas with more reef and greater distance from shore. Scarid abundance showed a negative correlation with protection and a positive correlation with distance to shore (Fig 6). In terms of body size of these families, the largest individuals were found inside the NTRs, with significant differences (Carangidae: *U* = 38283, p-value<0.001; Scaridae: *U* = 4462.0, p-value<0.001; Kyphosidae: *U* = 6450.5, p-value<0.001; Epinephelidae: *U* = 9341.5, p-value = 0.013) (Figs 6C, F and I).

**Fig 6.**
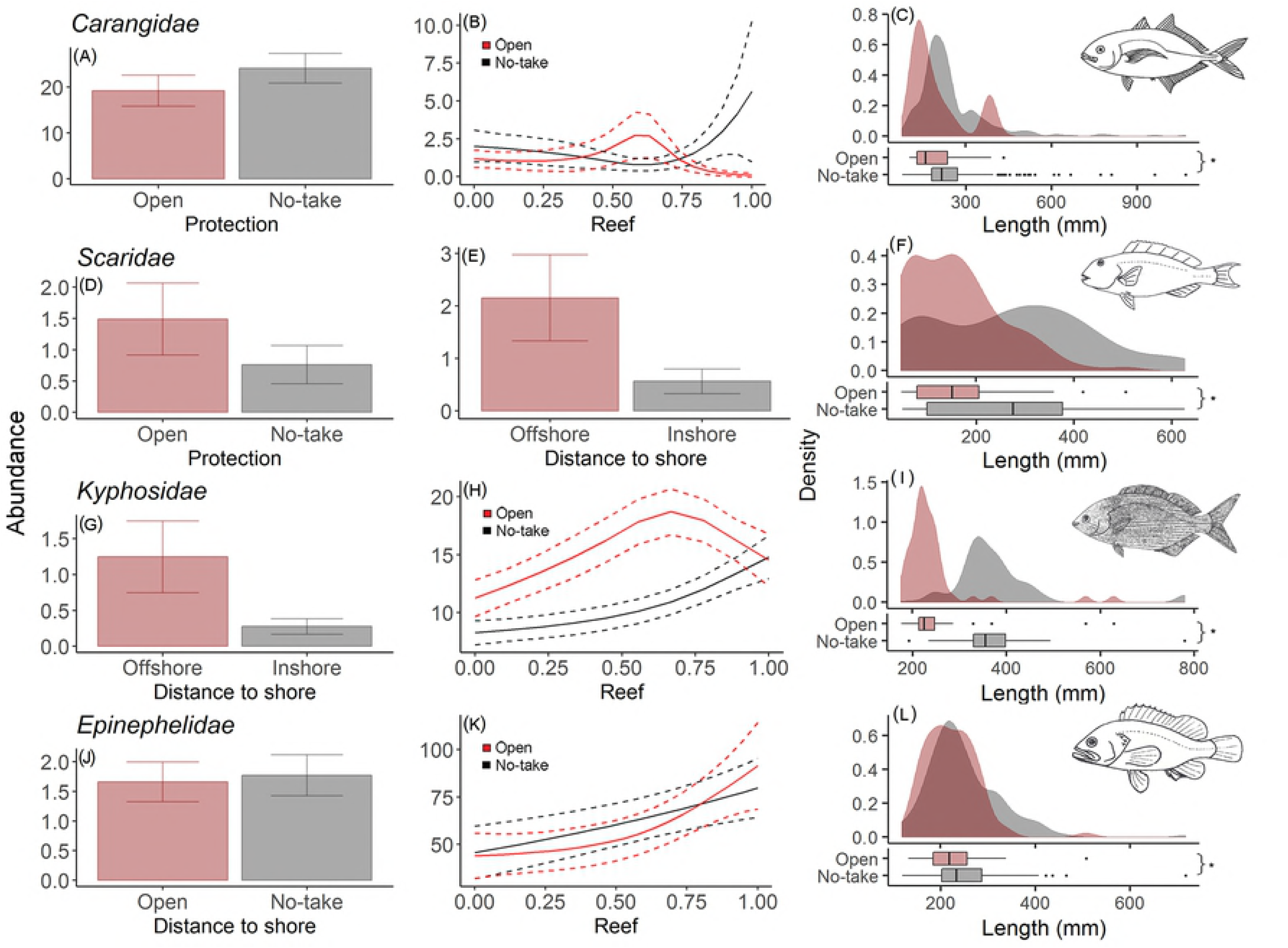
Plots of the most parsimonious models, Kernel density plots and boxplots for fork length (mm) for the most abundant and target families (A,B,C) Carangidae, (D,E,F) Scaridae, (G,H,I) Kyphosidae and (J,K,L) Epinephelidae. The dotted line represents 95% confidence interval. * Significant difference. Fish drawings (F, I and L) were based on Carvalho-Filho [50].

## Discussion

This study is the first to generate fisheries independent data using non-destructive stereo-video methods from the Southeastern Atlantic where limited information on fishing or fish ecology exists. Besides, the approach adopted here made it possible to distinguish the effect of fishing from habitat variables on different components of the fish assemblage, demonstrating how NTRs can be used as benchmarks to contribute to resource management and marine conservation.

### The role of no-take marine reserves

Broadly, total abundance and biomass were greater inside no-take areas, a pattern also registered in previous studies [5, 73, 74]. The assessment of biomass in the marine environment is relevant and can reveal the health status of an environment especially because it can be used to represent the energy flux, as well as the potential to provide ecosystem goods and services [75, 76]. Based on this, results indicate that the NTR in question is providing resources which are being effectively converted into biomass. Conversely, the opposite was found in areas open to fishing, presenting a decreased ecosystem functioning driven by the selective removal of large individuals [77, 78]. Higher overall abundance and biomass within NTRs indicates the significant removal of fish by fisheries in the open access areas in the region.

Distance from the coast was an important factor, explaining the higher richness, abundance and biomass recorded in islands further from the coast. This factor has demonstrated to influence fish assemblages structure in several coral and rocky reefs around the world [79– 82] and also in the Brazilian Province [11, 13, 23, 48]. The first hypotheses we raise to explain the higher richness and abundance in offshore islands may be related to the total area of rocky reefs. The reef in those islands reach a higher depth than coastal zones and the islands are larger, therefore presenting a larger continuous surface area of reef, in contrast with inshores coastlines that often present rocky shores interspersed with sandy beaches. Surface area of reef has been directly attributed to fish assemblage structure in some studies. For example, Francini-Filho and Moura [18] found a more pronounced increase of overall biomass over time in areas adjacent to coral reefs that reach deeper water. Furthermore, Roberts and Ormond [83] registered higher species richness with depth, and Gibran and Moura [48] also detected this tendency for rocky reefs in the same region of the present study. These findings might be due to higher availability of resources and a possible lower competition in offshore islands, especially for space [84].

The second hypothesis to explain the higher values of ecological metrics is related to the proximity of anthropogenic activities. The close proximity of human populations to a fish assemblage causes negative effects and is demonstrated worldwide [3, 77, 85–87]. Areas close to the mainland are easier to access and tend to have more fishing activities. Nearshore waters (<50 m water depth) of the São Paulo state coast, are highly explored by both artisanal and industrial fisheries fleets with artisanal, low mobility fleets most dominant in water depth <20 m [88]. Coastal regions with high population densities, such as São Paulo, are more exposed to human activities causing disturbances and changes in coastal dynamics, especially concerning the high input of nutrients and pollution through air deposition, river discharges, urban and industrial wastewater effluents, groundwater and surface runoff [89]. These potentially harmful components cause environmental stress and direct and indirectly cause damage to coastal biota [90]. In addition, areas near the coast also face greater exposure to major developments, such as harbors and marinas, which can also significantly change the coastal landscape, causing degradation of habitats and consequently affecting fish assemblage. Further studies in the region are needed to test these hypotheses in order to determine whether or how much of this pattern is explained by biogeographic traits or anthropogenic activities.

### Target and non-target species

Higher abundance of target species was observed within NTRs, but protection status did not correlate with any differences in the abundance of non-target species. Indeed, studies have showed increased abundance of highly targeted fishes inside no-take NTRs, with lower influence on non-target [3, 5, 22, 91, 92], reinforcing evidence of the direct effects of fishing. Abundance of target species also increased with greater distance from the shore, which can be related to the increased fisheries activity close to the shore as described above [88].

Conversely, species richness and richness of target and non-target fish, was not related to protection status, being mostly explained by relief. Higher species diversity in more complex environments have been described in the literature [3, 12], and is likely related to increased availability of food, decreased competition, and lower probability of predator-prey encounters [84, 93, 94]. Structurally complex environments have higher availability and diversity of niches, accommodating a higher number of species in a small area.

### Fish functional groups

Although functional groups responded differently to fisheries pressure, we found evidence that protection status affected the trophic structure of the fish assemblage, since carnivores/piscivores, mobile invertivore, and territorial herbivores were more abundant within the NTR, whilst sessile invertivores were less abundant. However, protection was not relevant for omnivores and roving herbivores. The carnivores/piscivores functional group is comprised of species targeted by fisheries in the region [16, 50–52] and is most likely the reason why their abundance was higher within NTRs. Even though relief was important, it was relevant only when combined with protection. These results suggests that the NTR is facilitating the recovery of top trophic group levels, which are usually the first group depleted by fisheries [78, 95, 96].

Some mobile invertivore species are targeted by fisheries, but are not considered as important to fisheries as carnivores because of their smaller body size, such as haemulids, labrids and small carangids. Nevertheless, protection was still an important factor to predict abundance of this group, suggesting some fisheries pressure, albeit less than highly targeted carnivores/piscivores. This might be related to a depletion of top predators, leading to an exploration of lower trophic levels, as already described worldwide [96], including Brazilian coast (Freire and Pauly, 2010) The other factor strongly affecting abundance of this group is distance to shore, which may be related to the larger rocky reef surface, offering more resources and, consequently, less competition [84]. This is especially important for small and benthic mobile invertivores of the families Blenniidae and Serranidae, that live closely associated with the substrate [50]. Similar results were found for omnivores, in which higher abundance is more likely to occur in high complex habitats in offshore islands, probably for the same reasons, since this group encompasses blennies, pomacentrids, pomacanthids and species of the order Tetraodontiformes. Although some species within this category are targeted by fisheries (Mugilidae, Sparidae, Ephippidae), protection was not an important factor to determine abundance. This is probably related to the plasticity of the omnivorous diet, which can enable greater resistance to environmental changes (e.g. [97, 98]).

As the abundance of sessile invertivores was very low in samples, the model was not robust. However, lower abundance found within NTRs and in more complex reefs indicated by the model may be related to the elusive behaviour of these species, which usually hide from divers and may not be recorded. Since these species feed on benthic invertebrates generally associated with hard substrate, we would expect a higher abundance in more topographic complex environments. For planktivores, models did not predict the abundance well, most likely because species in this group show highly variable body sizes, occupying very different niches. For example, fish from Echeneidae and Carangidae families are mobile and large-bodied species, occupying the pelagic environment, whilst the species from Pomacentridae and Pempheridae families are small-bodied species that live associated with burrows and crevices on the rocky reef [50]. Therefore, it was not possible to determine a single robust model to explain abundance of this functional group with the predictor variables used.

Abundance of roving herbivores was related to distance from shore and topographic complexity, which is expected considering its diet, algae and detritus, are mostly found in reef environments [99], which are more likely abundant in larger rocky reefs of offshore islands. This is similar with the results for territorial herbivores, in which protection was only important when combined with topographic complexity. This is also likely, since territorial herbivores, such as *Stegastes* ssp., are found in complex regions of the reef protecting colonies of the major components of their diet, primarily fast growing red and green filamentous algae [100, 101]. As habitat characteristics were more influential in herbivores abundance than protection status, fisheries effects were not evident for these groups, even though some of them are targeted in the region.

### Targeted fish families

One of the consequences of large removal of individuals by fisheries activities is represented by a rapid decrease in abundance and richness, especially of large bodied target species [78]. Indeed, the effects of fisheries on the size of individuals is well described, in which target species reach larger sizes within NTRs [22, 86, 91, 102, 103]. The present study corroborates these studies, showing a significantly higher density of larger individuals of target species of the families Epinephelidae, Kyphosidae, Carangidae and Scaridae within protected areas. This also represents an increase in reproduction capacity of these groups in protected areas as larger individuals usually present much higher fecundity. This increases the probability of exporting larvae from NTRs to adjacent areas [18, 104] repopulating fished reefs and helping to restock targeted species in fished areas.

Networks of moderate size (10-100 km^2^) NTRs have demonstrated to be more effective in resource management and conservation when compared to smaller protected areas [105]. However, small (1-5 km^2^) and very small (<1 km^2^) areas have been widely implemented and shown to have some advantages, specifically for small bodied and sedentary species with smaller home ranges [106–109]. In particular, individuals of the Epinephelidae family presented a higher abundance with protection and also in complex environments within the very small NTR in question. These species live associated with borrows within rocky reefs [110] and are highly targeted by fisheries, indicating that they may be the group benefiting most from protection, as seen in this study.

The abundance of kyphosids was not related to protection and was more abundant in regions offshore with the greater presence of reef. As this species is considered herbivorous, grazing predominately on macroalgae (*Sargassum* spp.) associated with rocks [111], we expect to record higher numbers at locations with greater food availability, including offshore areas with more rocky reef. However, larger individuals could be targeted by fishers, resulting in their higher abundance recorded within NTRs. This indicates that the NTRs are allowing the growth of individuals and, therefore, providing greater reproductive capacity for the species.

For the Carangidae family, an effect of protection in abundance was evident, suggesting a high removal, especially of large individuals, in areas open to fisheries. Besides, regardless of being a mobile species, they are frequently found associated with hard structures [35] and even following other species [112], and probably for this reason, individuals of this family have shown to benefit from NTRs in reefs [3, 113].

Fish of the Scaridae family showed a higher abundance in fished areas, likely due to the absence of top predators (carnivores/piscivores), since species of this family have been registered to be preyed upon by epinephelids, carangids and muraenids [114]. Even though they were more abundant in fished areas, fish size was smaller, representing a fishing pressure in larger sizes, as also described by Floeter et al. [22]. Also, the abundance of these roving herbivores was higher with distance from the coast, what could be related to the availability of food and lower competition in larger and continuous reefs offshore.

## Conclusions

Rocky reefs in Brazil do not have high species richness of fish when compared to reefs worldwide, however, these areas stand out for the proportion of endemic and endangered species concentrated in small areas [25, 26, 115]. Therefore, it is crucial to generate information about the role NTRs can play in fish assemblages of this region. In addition, a better understanding of patterns in the effects of fishing on a fish assemblage provides robust metrics for conservation and fisheries management, whilst also providing information on focal species and biological variables most relevant to monitor this effectiveness.

The present study supports other findings adding empirical evidence that very small NTRs (<1km^2^) can protect fish assemblages from the direct effects of fishing, increasing abundance and biomass, especially of targeted species, therefore contributing to the management of fisheries resources at a local and regional scale. Some functional groups tended to benefit more from protection, such as carnivores/piscivores and mobile invertivores, while others tended to decrease in abundance, such as the Scaridae family. Another outstanding difference is concerning body size, mostly for target species, in which NTRs allow target species to reach larger sizes. The Epinephelidae family showed greater evidence to benefit from these very small NTRs, especially due to its high importance to fisheries and its small home range. However, we recommend that fully representative and replicated networks of larger NTRs (>10km2) should be established in the region and will provide a much more robust framework for investigating the effects of fishing and informing conservation and fisheries management more broadly.

As a concluding remark, our findings show strong influence of protection, distance from the shore and mean relief on fish assemblage characteristics, in which protected areas further from the human influence and with a higher topographic complexity tend to have greater abundance and biomass of fish. Our results highlight the crucial role these areas play in the maintenance and repopulation of highly valuable commercial species to the fishing activity of the region, displaying the importance of keeping and implementing other small NTRs in the region. The use of stereo-videos in this study has shown to be effective and feasible in this region, providing valuable and robust information to aid conservation and fisheries management in Brazil.

## Acknowledgments

The authors would like to acknowledge the Chico Mendes Institute for Biodiversity Conservation (ICMBio) for the permits (#48259-1) and field support and also to the Universo Marinho Dive Company and the staff for sampling assistance and equipment. We are also thankful to Instituto Biopesca for all the funding management, Alfredo Carvalho Filho for the fish images and the anonymous reviewers for improving the paper.

## Supporting Information

S1 Table List of species found within no-take reserves and fished areas. *Endemic from Brazilian biogeographic province; VU_I_= Vulnerable by International Union for Nature Protection (IUCN) Red List; NT_I_= Near threatened by IUCN; VU_Br_= Vulnerable by Brazilian legislation [72]; CR_Br_= Critically endangered by Brazilian Legislation; Y= Target; N= Non-target; N= Abundance; C%= Commonality.

## References

1. Lester S, Halpern B, Grorud-Colvert K, Lubchenco J, Ruttenberg B, Gaines S, et al. Biological effects within no-take marine reserves: a global synthesis. Mar Ecol Prog Ser. 2009;384: 33–46.

2. Gell FR, Roberts CM. The fishery effects of marine reserves and fishery closures. Washington, DC.: World Wildlife Fund; 2002.

3. Edgar GJ, Stuart-Smith RD, Willis TJ, Kininmonth S, Baker SC, Banks S, et al. Global conservation outcomes depend on marine protected areas with five key features. Nature. 2014;506: 216–220.

4. Worm B, Barbier EB, Beaumont N, Duffy JE, Folke C, Halpern BS, et al. Impacts of biodiversity loss on ocean ecosystem services. Science. 2006;314: 787–790.

5. Micheli F, Halpern BS, Botsford LW, Warner RR. Trajectories and correlates of community change in no-take marine reserves. Ecol Appl. 2004;14: 1709–1723.

6. Steneck RS. Human influences on coastal ecosystems: does overfishing create trophic cascades? Trends Ecol Evol. 1998;13: 429–430.

7. Graham NAJ, Evans RD, Russ GR. The effects of marine reserve protection on the trophic relationships of reef fishes on the Great Barrier Reef. Environ Conserv. Cambridge University Press; 2003;30: 200–208.

8. Krebs CJ. Ecology: The experimental analysis of distribution and abundance. New York: Harper & Row; 1972.

9. Tokeshi M, Arakaki S. Habitat complexity in aquatic systems: fractals and beyond. Hydrobiologia. 2012;685: 27–47.

10. Langlois TJ, Radford BT, Van Niel KP, Meeuwig JJ, Pearce AF, Rousseaux CSG, et al. Consistent abundance distributions of marine fishes in an old, climatically buffered, infertile seascape: Abundance distributions of fishes in stable seascapes. Glob Ecol Biogeogr. 09/2012;21: 886–897.

11. Teixeira-Neves TP, Neves LM, Araújo FG. Hierarchizing biological, physical and anthropogenic factors influencing the structure of fish assemblages along tropical rocky shores in Brazil. Environ Biol Fishes. 2015;98: 1645–1657.

12. Neves LM, Teixeira-Neves TP, Pereira-Filho GH, Araújo FG. The farther the better: effects of multiple environmental variables on reef fish assemblages along a distance gradient from river influences. PLoS One. 2016;11: e0166679.

13. Morais RA, Ferreira CEL, Floeter SR. Spatial patterns of fish standing biomass across Brazilian reefs. J Fish Biol. 2017; doi:10.1111/jfb.13482

14. García-Charton JA, Pérez-Ruzafa Á, Sánchez-Jerez P, Bayle-Sempere JT, Reñones O, Moreno D. Multi-scale spatial heterogeneity, habitat structure, and the effect of marine reserves on Western Mediterranean rocky reef fish assemblages. Mar Biol. Springer-Verlag; 2004;144: 161–182.

15. Parsons DF, Suthers IM, Cruz DO, Smith JA. Effects of habitat on fish abundance and species composition on temperate rocky reefs. Mar Ecol Prog Ser. 2016;561: 155–171.

16. Floeter SR, Krohling W, Gasparini JL, Ferreira CEL, Zalmon IR. Reef fish community structure on coastal islands of the southeastern Brazil: the influence of exposure and benthic cover. Environ Biol Fishes. 2007;78: 147–160.

17. ICMBio. Cadastro Nacional de Unidades de Conservação. In: Instituto Chico Mendes de Proteção à Natureza [Internet]. 7 Jan 2018 [cited 29 Aug 2018]. Available: http://www.mma.gov.br/areas-protegidas/cadastro-nacional-de-ucs/dados-consolidados.html

18. Francini-Filho RB, Moura RL. Dynamics of fish assemblages on coral reefs subjected to different management regimes in the Abrolhos Bank, eastern Brazil. Aquat Conserv. 2008;18: 1166–1179.

19. Anderson A, Bonaldo R, Barneche D, Hackradt C, Félix-Hackradt F, García-Charton J, et al. Recovery of grouper assemblages indicates effectiveness of a marine protected area in Southern Brazil. Mar Ecol Prog Ser. 2014;514: 207–215.

20. Anderson AB, Batista MB, Gibran FZ, Félix-Hackradt FC, Hackradt CW, García-Charton JA, et al. Habitat use of five key species of reef fish in rocky reef systems of southern Brazil: evidences of MPA effectiveness. Mar Biodivers. 2018; doi:10.1007/s12526-018-0893-6

21. Ilarri MI, Souza AT, Rosa RS. Community structure of reef fishes in shallow waters of the Fernando de Noronha archipelago: effects of different levels of environmental protection. Mar Freshwater Res. CSIRO PUBLISHING; 2017;68: 1303–1316.

22. Floeter SR, Halpern BS, Ferreira CEL. Effects of fishing and protection on Brazilian reef fishes. Biol Conserv. 2005;128: 391–402.

23. Floeter SR, Guimaraes RZP, Rocha LA, Ferreira CEL, Rangel CA, Gasparini JL. Geographic variation in reef-fish assemblages along the Brazilian coast. Glob Ecol Biogeogr. Wiley Online Library; 2001;10: 423–431.

24. Rocha LA. Patterns of distribution and processes of speciation in Brazilian reef fishes. J Biogeogr. Blackwell Science Ltd; 2003;30: 1161–1171.

25. Moura RL. Brazilian reefs as priority areas for biodiversity conservation in the Atlantic Ocean. Proceeding of the 9th International Coral Reef Symposium, Bali, Indonesia. researchgate.net; 2002. pp. 917–920.

26. Pinheiro HT, Rocha LA, Macieira RM, Carvalho-Filho A, Anderson AB, Bender MG, et al. South-western Atlantic reef fishes: Zoogeographical patterns and ecological drivers reveal a secondary biodiversity centre in the Atlantic Ocean. Treml E, editor. Divers Distrib. 2018;24: 951–965.

27. Aued AW, Smith F, Quimbayo JP, Cândido DV, Longo GO, Ferreira CEL, et al. Large-scale patterns of benthic marine communities in the Brazilian Province. PLoS One. 2018;13: e0198452.

28. Thompson AA, Mapstone BD. Observer effects and training in underwater visual surveys of reef fishes. Mar Ecol Prog Ser. 1997;154: 53–63.

29. St. John J, Russ GR, Gladstone W. Accuracy and bias of visual estimates of numbers, size structure and biomass of a coral reef fish. Mar Ecol Prog Ser. Inter-Research Science Center; 1990;64: 253–262.

30. Harvey ES, Fletcher D, Shortis M. A comparison of the precision and accuracy of reef fish length divers and stereo. Fish Bull. 2000;99: 63–71.

31. Harvey E, Fletcher D, Shortis M, Kendrick G. A comparison of underwater visual distance estimates made by scuba divers and a stereo-video system: implications for underwater visual census of reef fish abundance. Mar Freshwater Res. 2004;55: 573–580.

32. Cappo M, Harvey E, Shortis M. Counting and measuring fish with baited video techniques—an overview. In: Lyle JM, Furlani DM, Buxton CD, editors. Australian Society for Fish Biology Workshop Proceedings. Tasmania: Australian Society for Fish Biology; 2006. pp. 101–114.

33. Goetze JS, Jupiter SD, Langlois TJ, Wilson SK, Harvey ES, Bond T, et al. Diver operated video most accurately detects the impacts of fishing within periodically harvested closures. J Exp Mar Bio Ecol. 2015;462: 74–82.

34. Whitmarsh SK, Fairweather PG, Huveneers C. What is Big BRUVver up to? Methods and uses of baited underwater video. Rev Fish Biol Fish. Springer International Publishing; 2017;27: 53–73.

35. Bond T, Partridge JC, Taylor MD, Langlois TJ, Malseed BE, Smith LD, et al. Fish associated with a subsea pipeline and adjacent seafloor of the North West Shelf of Western Australia. Mar Environ Res. 2018; doi:10.1016/j.marenvres.2018.08.003

36. Harvey E, Cappo M, Butler J, Hall N, Kendrick G. Bait attraction affects the performance of remote underwater video stations in assessment of demersal fish community structure. Mar Ecol Prog Ser. 2007;350: 245–254.

37. Langlois TJ, Newman SJ, Cappo M, Harvey ES, Rome BM, Skepper CL, et al. Length selectivity of commercial fish traps assessed from in situ comparisons with stereo-video: Is there evidence of sampling bias? Fish Res. 2015;161: 145–155.

38. Holmes TH, Wilson SK, Travers MJ, Langlois TJ, Evans RD, Moore GI, et al. A comparison of visual- and stereo-video based fish community assessment methods in tropical and temperate marine waters of Western Australia. Limnol Oceanogr Methods. 2013;11: 337–350.

39. Watson DL, Harvey ES, Anderson MJ, Kendrick GA. A comparison of temperate reef fish assemblages recorded by three underwater stereo-video techniques. Mar Biol. 2005;148: 415–425.

40. Brasil. Decreto No 94.656, de 20 de Julho de 1987 [Internet]. 1987 [cited 29 Aug 2018]. Available: http://www.planalto.gov.br/ccivil_03/decreto/1980-1989/1985-1987/D94656.htm

41. Emplasa. Empresa Paulista de Planejamento Metropolitano. In: Ortofotos do Estado de São Paulo - 2010/2011. Projeto Mapeia São Paulo. Infraestrutura de Dados Espaciais Ambientais do Estado de São Paulo – IDEA-SP. Secretaria do Meio Ambiente do Estado de São Paulo [Internet]. 1 Jan 2010 [cited 13 Sep 2018]. Available: datageo.ambiente.sp.gov.br

42. CFA/DPM. Coordenadoria de Fiscalização Ambiental. In: Mapa da Pesca Sustentável. Infraestrutura de Dados Espaciais Ambientais do Estado de São Paulo – IDEA-SP. Secretaria do Meio Ambiente do Estado de São Paulo. [Internet]. 31 Dec 2014 [cited 13 Sep 2018]. Available: datageo.ambiente.sp.gov.br

43. Dorman SR, Harvey ES, Newman SJ. Bait effects in sampling coral reef fish assemblages with stereo-BRUVs. PLoS One. 2012;7: e41538.

44. Whitelaw AW, Sainsbury KJ, Dews GJ, Campbell RA. Catching characteristics of four fish-trap types on the North West Shelf of Australia. Mar Freshwater Res. CSIRO PUBLISHING; 1991;42: 369–382.

45. Langlois T, Williams J, Monk J, Bouchet P, Currey L, Goetze J, et al. Marine sampling field manual for benthic stereo-BRUVS (Baited Remote Underwater Videos). In: Przeslawski R FS, editor. Field Manuals for Marine Sampling to Monitor Australian Waters. National Environmental Science Programme (NESP); 2018. pp. 82–104.

46. Langlois TJ, Friedman A, Goetze J, Bond T, Birt MJ, McLean DL, et al. GlobalArchive: a repository of aquatic annotation. Google Docs. 2017; 18.

47. Froese R, Pauly D. FishBase. Version (06/2018) [Internet]. 2018 [cited 27 Aug 2018]. Available: www.fishbase.org

48. Gibran FZ, Moura RL de. The structure of rocky reef fish assemblages across a nearshore to coastal islands’ gradient in Southeastern Brazil. Neotrop Ichthyol. 2012;10: 369–382.

49. Ferreira CEL, Gonçalves JEA, Coutinho R. Community structure of fishes and habitat complexity on a tropical rocky shore. Environ Biol Fishes. Kluwer Academic Publishers; 2001;61: 353–369.

50. Carvalho-Filho A. Peixes: Costa brasileira. São Paulo: Editora Melro; 1999.

51. Rolim FA, Rodrigues PFC, Gadig OBF. Peixes de recife rochoso: Estação Ecológica de Tupinambás. São Paulo: Anolis Books; 2017.

52. Begossi A, Richerson PJ. Biodiversity, family income and ecological niche: A study on the consumption of animal foods on Búzios Island (Brazil). Ecol Food Nutr. Routledge; 1993;30: 51–61.

53. Bennett K, Wilson SK, Shedrawi G, McLean DL, Langlois TJ. Can diver operated stereo-video surveys for fish be used to collect meaningful data on benthic coral reef communities?: Comparing benthos from video methods. Limnol Oceanogr Methods. 2016;14: 874–885.

54. Cappo M, Stowar M, Syms C, Johansson C, Cooper T. Fish-habitat associations in the region offshore from James Price Point-a rapid assessment using Baited Remote Underwater Video Stations (BRUVS). J R Soc West Aust. 2011;94: 303–321.

55. Wilson SK, Babcock RC, Fisher R, Holmes TH, Moore JAY, Thomson DP. Relative and combined effects of habitat and fishing on reef fish communities across a limited fishing gradient at Ningaloo. Mar Environ Res. 2012;81: 1–11.

56. Althaus F, Hill N, Ferrari R, Edwards L, Przeslawski R, Schönberg CHL, et al. A standardised vocabulary for identifying benthic biota and substrata from underwater imagery: The CATAMI Classification Scheme. PLoS One. 2015;10: e0141039.

57. Wilson SK, Graham NAJ, Polunin NVC. Appraisal of visual assessments of habitat complexity and benthic composition on coral reefs. Mar Biol. 2007;151: 1069–1076.

58. Lin X, Zhang D. Inference in generalized additive mixed models by using smoothing splines. J R Stat Soc Series B Stat Methodol. 1999;61: 381–400.

59. Hastie T, Tibshirani R. Generalized Additive Models. Stat Sci. Institute of Mathematical Statistics; 1986;1: 297–310.

60. Fisher R, Wilson SK, Sin TM, Lee AC, Langlois TJ. A simple function for full-subsets multiple regression in ecology with R. Ecol Evol. 2018; doi:10.1002/ece3.4134

61. Akaike H. Information theory and an extension of the maximum likelihood principle. In: BN Petrov; F Csaki, editor. 2nd International Symposium on Information Theory. Budapest, Hungary: Akadémiai Kiadó; 1973. pp. 267–281.

62. Burnham KP, Anderson DR. Model selection and multimodel Inference: A practical information-theoretic approach. 2nd Edition. New York: Springer Science & Business Media; 2003.

63. R Core Team. R Language for Statistical Computing. In: R: A Language and Environment for Statistical Computing [Internet]. Vienna, Austria: R Foundation for Statistical Computing; 2018 [cited 8 Aug 2018]. Available: https://www.R-project.org/

64. Wood S, Scheipl F. gamm4: Generalized Additive Mixed Models using “mgcv” and “lme4” [Internet]. 2017 [cited 27 Aug 2018]. Available: https://CRAN.R-project.org/package=gamm4

65. Wood SN. Generalized additive models: An introduction with R. Boca Raton, FL: CRC Press; 2006.

66. Barton K. MuMIn: Multi-Model Inference [Internet]. 2018 [cited 27 Aug 2018]. Available: https://CRAN.R-project.org/package=MuMIn

67. Microsoft-Corporation, Weston S. doParallel: Foreach Parallel Adaptor for the “parallel” Package [Internet]. 2017. Available: https://CRAN.R-project.org/package=doParallel

68. Wickham H, François R, Henry L, Müller K. dplyr: A Grammar of Data Manipulation [Internet]. 2018 [cited 27 Aug 2018]. Available: https://CRAN.R-project.org/package=dplyr

69. Floeter SR, Gasparini JL. The southwestern Atlantic reef fish fauna: composition and zoogeographic patterns. J Fish Biol. Wiley Online Library; 2000;56: 1099–1114.

70. Joyeux JC, Floeter SR, Ferreira CEL, Gasparini JL. Biogeography of tropical reef fishes: the South Atlantic puzzle. J Biogeogr. Wiley; 2001;28: 831–841.

71. Gasparini JL, Joyeux J-C, Floeter SR. Sparisoma tuiupiranga, a new species of parrotfish (Perciformes: Labroidei: Scaridae) from Brazil, with comments on the evolution of the genus. Zootaxa. 2003;384: 1–14.

72. MMA. Portaria MMA No 445, de 17 de Dezembro de 2014. In: Ministério do Meio Ambiente, Brasil [Internet]. 2014. Available: http://www.icmbio.gov.br/cepsul/images/stories/legislacao/Portaria/2014/p_mma_445_2014_lista_peixes_amea%C3%A7ados_extin%C3%A7%C3%A3o.pdf

73. Soler GA, Edgar GJ, Thomson RJ, Kininmonth S, Campbell SJ, Dawson TP, et al. Reef fishes at all trophic levels respond positively to effective marine protected areas. PLoS One. 2015;10: e0140270.

74. Halpern BS. The impact of marine reserves: do reserves work and does reserve size matter? Ecol Appl. 2003;13: 117–137.

75. Cardinale BJ, Srivastava DS, Emmett Duffy J, Wright JP, Downing AL, Sankaran M, et al. Effects of biodiversity on the functioning of trophic groups and ecosystems. Nature. Nature Publishing Group; 2006;443: 989.

76. Hooper DU, Chapin FS III, Ewel JJ, Hector A, Inchausti P, Lavorel S, et al. Effects of biodiversity on ecosystem functioning: a consensus of current knowledge. Ecol Monogr. 2005;75: 3–35.

77. Mora C, Aburto-Oropeza O, Bocos AA, Ayotte PM, Banks S, Bauman AG, et al. Global human footprint on the linkage between biodiversity and ecosystem functioning in reef fishes. PLoS Biol. Public Library of Science; 2011;9: e1000606.

78. Crowder LB, Hazen EL, Avissar N, Bjorkland R, Latanich C, Ogburn MB. The impacts of fisheries on marine ecosystems and the transition to ecosystem-based management. Annu Rev Ecol Evol Syst. 2008;39: 259–278.

79. Van Nguyen L, Kim Phan H. Distribution and factors influencing on structure of reef fish communities in Nha Trang Bay Marine Protected Area, South-Central Vietnam. Environ Biol Fishes. 2007;82: 309.

80. Lecchini D, Adjeroud M, Pratchett MS, Cadoret L, Galzin R. Spatial structure of coral reef fish communities in the Ryukyu Islands, southern Japan. Oceanol Acta. 2003;26: 537–547.

81. Schultz AL, Malcolm HA, Bucher DJ, Linklater M, Smith SDA. Depth and medium-scale spatial processes influence fish assemblage structure of unconsolidated habitats in a subtropical marine park. PLoS One. journals.plos.org; 2014;9: e96798.

82. Malcolm HA, Jordan A, Smith SDA. Biogeographical and cross-shelf patterns of reef fish assemblages in a transition zone. Mar Biodivers. 2010;40: 181–193.

83. Roberts CM, Ormond R. Habitat complexity and coral reef fish diversity and abundance on Red Sea fringing reefs. Mar Ecol Prog Ser. 1987;41: 1–8.

84. Sale PF. Maintenance of high diversity in coral reef fish communities. Am Nat. [University of Chicago Press, American Society of Naturalists]; 1977;111: 337–359.

85. Babcock RC, Shears NT, Alcala AC, Barrett NS, Edgar GJ, Lafferty KD, et al. Decadal trends in marine reserves reveal differential rates of change in direct and indirect effects. Proc Natl Acad Sci U S A. 2010;107: 18256–18261.

86. Edgar GJ, Stuart-Smith RD, Thomson RJ, Freeman DJ. Consistent multi-level trophic effects of marine reserve protection across northern New Zealand. PLoS One. 2017;12: 12. e0177216.

87. Langlois TJ, Harvey ES, Meeuwig JJ. Strong direct and inconsistent indirect effects of fishing found using stereo-video: Testing indicators from fisheries closures. Ecol Indic. Elsevier; 2012;23: 524–534.

88. Imoto RD, Carneiro MH, Ávila-da-Silva AO. Spatial patterns of fishing fleets on the Southeastern Brazilian Bight. Lat Am J Aquat Res. 2016;44. doi:10.3856/vol4-issue-fulltext-12

89. von Glasow R, Jickells TD, Baklanov A, Carmichael GR, Church TM, Gallardo L, et al. Megacities and large urban agglomerations in the coastal zone: interactions between atmosphere, land, and marine ecosystems. Ambio. 2013;42: 13–28.

90. Adams SM. Assessing cause and effect of multiple stressors on marine systems. Mar Pollut Bull. 2005;51: 649–657.

91. Malcolm HA, Williams J, Schultz AL, Neilson J, Johnstone N, Knott NA, et al. Targeted fishes are larger and more abundant in “no-take” areas in a subtropical marine park. Estuar Coast Shelf Sci. Elsevier; 2018;212: 118–127.

92. Evans RD, Russ GR. Larger biomass of targeted reef fish in no-take marine reserves on the Great Barrier Reef, Australia. Aquat Conserv. Wiley Online Library; 2004;14: 505–519.

93. Eklöv P, Diehl S. Piscivore efficiency and refuging prey: the importance of predator search mode. Oecologia. 1994;98: 344–353.

94. Hauzy C, Tully T, Spataro T, Paul G, Arditi R. Spatial heterogeneity and functional response: an experiment in microcosms with varying obstacle densities. Oecologia. 2010;163: 625–636.

95. Christensen V, Guenette S, Heymans JJ, Walters CJ, Watson R, Zeller D, et al. Hundred-year decline of North Atlantic predatory fishes. Fish Fish. 2003;4: 1–24.

96. Pauly D, Christensen V V, Dalsgaard J, Froese R, Torres F Jr. Fishing down marine food webs. Science. science.sciencemag.org; 1998;279: 860–863.

97. Bellwood DR, Hoey AS, Ackerman JL, Depczynski M. Coral bleaching, reef fish community phase shifts and the resilience of coral reefs. Glob Chang Biol. 2006;12: 1587–1594.

98. Pratchett MS, Hoey AS, Wilson SK, Messmer V, Graham NAJ. Changes in biodiversity and functioning of reef fish assemblages following coral bleaching and coral loss. Diversity. Molecular Diversity Preservation International; 2011;3: 424–452.

99. Francini-Filho RB, Ferreira CM, Coni EOC, De Moura RL, Kaufman L. Foraging activity of roving herbivorous reef fish (Acanthuridae and Scaridae) in eastern Brazil: influence of resource availability and interference competition. J Mar Biol Assoc U K. Cambridge University Press; 2010;90: 481–492.

100. Ferreira CEL, Gonçalves JEA, Coutinho R, Peret AC. Herbivory by the Dusky Damselfish Stegastes fuscus (Cuvier, 1830) in a tropical rocky shore: effects on the benthic community. J Exp Mar Bio Ecol. 1998;229: 241–264.

101. Ferreira CM, Coni EOC, Medeiros DV, Sampaio CLS, Reis-Filho JA, Barros F, et al. Community structure of shallow rocky shore fish in a tropical bay of the southwestern Atlantic. Brazil J Oceanogr. Instituto Oceanográfico da Universidade de São Paulo; 2015;63: 379–396.

102. Watson DL, Anderson MJ, Kendrick GA, Nardi K, Harvey ES. Effects of protection from fishing on the lengths of targeted and non-targeted fish species at the Houtman Abrolhos Islands, Western Australia. Mar Ecol Prog Ser. 2009;384: 241–249.

103. DeMartini EE, Friedlander AM, Sandin SA, Sala E. Differences in fish-assemblage structure between fished and unfished atolls in the northern Line Islands, central Pacific. Mar Ecol Prog Ser. 2008;365: 199–215.

104. Palumbi SR. Marine reserves and ocean neighborhoods: The spatial scale of marine populations and their management. Annu Rev Environ Resour. 2004;29: 31–68.

105. Halpern BS, Warner RR. Matching marine reserve design to reserve objectives. Proc Biol Sci. 2003;270: 1871–1878.

106. Bonaldo RM, Pires MM, Guimarães PR Junior, Hoey AS, Hay ME. Small marine protected areas in Fiji provide refuge for reef fish assemblages, feeding groups, and corals. PLoS One. 2017;12: e0170638.

107. Di Franco A, Plass-Johnson JG, Di Lorenzo M, Meola B, Claudet J, Gaines SD, et al. Linking home ranges to protected area size: The case study of the Mediterranean Sea. Biol Conserv. 2018;221: 175–181.

108. Afonso P, Fontes J, Santos RS. Small marine reserves can offer long term protection to an endangered fish. Biol Conserv. 2011;144: 2739–2744.

109. McLaren BW, Langlois TJ, Harvey ES, Shortland-Jones H, Stevens R. A small no-take marine sanctuary provides consistent protection for small-bodied by-catch species, but not for large-bodied, high-risk species. J Exp Mar Bio Ecol. 2015;471: 153–163.

110. Gibran FZ. Activity, habitat use, feeding behavior, and diet of four sympatric species of Serranidae (Actinopterygii: Perciformes) in southeastern Brazil. Neotrop Ichthyol. Sociedade Brasileira de Ictiologia; 2007;5: 387–398.

111. Silvano RAM, Güth AZ. Diet and feeding behavior of Kyphosus spp. (Kyphosidae) in a Brazilian subtropical reef. Braz Arch Biol Technol. Tecpar; 2006;49: 623–629.

112. Sazima C, Krajewski JP, Bonaldo RM, Sazima I. Nuclear-follower foraging associations of reef fishes and other animals at an oceanic archipelago. Environ Biol Fishes. 2007;80: 351–361.

113. Santana-Garcon J, Newman SJ, Langlois TJ, Harvey ES. Effects of a spatial closure on highly mobile fish species: an assessment using pelagic stereo-BRUVs. J Exp Mar Bio Ecol. 2014;460: 153–161.

114. Randall JE. Food habits of reef fishes of the West Indies. Institute of Marine Sciences, University of Miami Coral Gables; 1967; Available: http://www.aoml.noaa.gov/general/lib/CREWS/Cleo/PuertoRico/prpdfs/randall-habits.pdf

115. Floeter SR, Rocha LA, Robertson DR, Joyeux JC, Smith-Vaniz WF, Wirtz P, et al. Atlantic reef fish biogeography and evolution. J Biogeogr. 2007;35: 22–47.

